# A general mechanism of air-borne hearing in Recent and early non-tympanate tetrapods

**DOI:** 10.1101/2022.05.09.491176

**Authors:** Jakob Christensen-Dalsgaard, Tanya Bojesen Lauridsen, Grace Capshaw, Catherine E. Carr

## Abstract

Tetrapod tympanic hearing probably emerged in the Triassic with independent origins in each of the major groups, more than 120 Myr after the water-land transition. During this long period, any auditory sensitivity must have been based on non-tympanic, bone conduction mechanisms. However, ‘bone conduction’ is a non-specific term describing several different modes of vibration that can stimulate the inner ear.

To understand hearing in a non-tympanic ear, we focus on the simplest model: that sound translates the head, i.e., that the head is pushed and pulled by the sound wave, and that this vibration is transduced by the inner ear. Simple translation is the mode of human low-frequency bone conduction sensitivity and translation by underwater sound is also the mode of auditory stimulation for most fishes. It is therefore a straightforward assumption that this may have been the mechanism of hearing in the early tetrapods. According to acoustic theory, the efficiency of translation of an object by sound is determined by its density and ka, the product of the acoustic wavenumber (k) and the radius (a) of the head. Simple finite-element models of translation by sound show that vibration velocities only depend on ka and density (for objects of the same shape and composition) and are almost constant (between 4 and 5 µm/s/Pa depending on shape) for objects with ka<1. We compare sensitivity to sound and to vibrations of the skull in animals lacking tympanic middle ears (snakes, salamanders, earless frogs, and lungfish) and show that the low-frequency air-borne sound sensitivity in these species is largely consistent with a translation mechanism. How translation of the head or body can stimulate the inner ear is most evident in an inertial system like the otolithic/otoconial ears of fish and early tetrapods, but fluid inertia in the inner ear may also generate hydrodynamic waves that can stimulate hair cells in the tetrapod inner ear, providing a mechanism for this simple mode of sound reception to confer hearing in earless animals.

## Introduction

The tympanic ear is described as an impedance matching device that counteracts the reflection of sound that would otherwise be caused by the large impedance difference between the air medium and animal tissue (roughly with the same impedance as water). More accurately, middle ear function is described as impedance transformation, since sound energy is collected and transformed into another form of mechanical energy (Christensen-Dalsgaard and Manley, 2013) that, in a sensitive middle ear, will generate high-amplitude vibrations at the oval window. The effects of this transformation are large; in humans with a non-functional middle ear, conductive hearing loss results in a decrease in sensitivity by up to 50 dB (Feldman 1963). Thus, the tympanic middle ear of tetrapods is often seen as a prerequisite for sensitive hearing in air.

However, tympanic middle ears emerged late in tetrapod evolution – in the Triassic, approximately 120 Myr after the origin of tetrapods and independently in all the major groups (Clack 1997, Christensen-Dalsgaard and Carr, 2008). Many Recent tetrapod species (snakes, some lizards, and many amphibians) have secondarily lost the functional middle ear system and the mechanism of their air-borne hearing is as unclear as it is in the early tetrapods. For example, 38 instances of secondary loss of middle ear functionality have been described in anuran amphibians (in the so-called ‘earless’ frogs, (Jaslow et al., 1988)), many of these in the bufonid family (Pereyra et al., 2016). A comparison of the sensitivity of earless toads with similar-sized tympanate species showed comparable sensitivity below 900 Hz, but thresholds to higher frequencies in ‘eared’ species were up to 20 dB lower. It has been known for some time that anurans possess ‘extratympanic’ sensitivity that dominates auditory sensitivity at low frequencies (below 400 Hz), where the eardrum shows very little response to sound (Jørgensen and Christensen-Dalsgaard, 1997a; Jørgensen and Christensen-Dalsgaard, 1997b; Lombard and Straughan, 1974; Wilczynski et al., 1987, as reviewed by Christensen-Dalsgaard (2005). Also, several studies have shown that the low-frequency auditory fibers are very sensitive to substrate vibrations (Christensen-Dalsgaard and Jørgensen, 1996; Christensen-Dalsgaard and Narins, 1993; Jørgensen and Christensen-Dalsgaard, 1991; Yu et al., 1991). Studies of air-borne hearing in atympanate snakes (Christensen et al., 2012) and salamanders (Capshaw et al., 2020) have shown similar sensitivities to low-frequency sounds (recent review in Capshaw et al. 2022).

In animals that lack tympanic middle ears to transform airborne sound into mechanical vibrations, hearing may occur as the detection of sound-generated translation of the head or body. This bone conduction mechanism is phylogenetically widespread, observed in lungfish, salamanders, and snakes, and therefore may represent a common pathway for sound detection in the absence of a functional middle ear. Here we evaluate a simple finite-element model of object translation by sound energy, and we compare this model to empirical findings in both tympanate and atympanate vertebrate species. We propose that, in animals for which the acoustical size of the receiver (the head) is small, translation by sound may be the primary mode for transferring airborne acoustic energy to the inner ear.

## Sound as a mechanical force -– underwater sound

### Box 1.

The interaction of an animal with sound depends on

1. the relationship between wavenumber (k) and size (radius, a) of the animal, expressed as

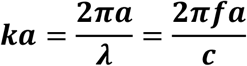

where *c* is speed of sound, *λ* wavelength, and *f* frequency, and
2. the difference between the density of the medium and the density of the animal.

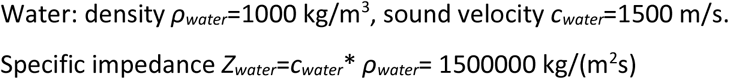

If this difference is very small, as for an animal in water, the animal will move with the medium particle velocity, *v*_*p*_=*p*/*Z*_*water*_, where *p* is sound pressure. For an underwater sound pressure of 1 Pa, *v*_*p*_ is 1/1500000 m/s = 0.6 µm/s. This particle velocity corresponds to a displacement of *d*=*v*_*p*_*/2πf*.

Underwater sounds relevant to most aquatic non-mammalian vertebrates tend to be below 5 kHz, therefore the wavelength is usually much larger than the animal and the animal’s acoustical size, ka, is less than 1. Perhaps the simplest form of interaction between an object and sound occurs when the sound generates a translational movement in the animal with the same particle velocity (see box 1) as the surrounding medium. Since the density of the animal is roughly equal to the density of the surrounding water, the animal has nearly neutral buoyancy and will be pushed and pulled by the sound pressure. In fish, the inner ear sensory cells are stimulated by inertial movement of the comparatively dense otoliths relative to this body motion. The particle velocity amplitude for a 1 Pa sound is 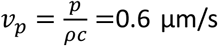 (box 1). This particle velocity at 1 Pa corresponds to displacements in the nanometer range for most frequencies, which is close to hair cell thresholds, and suggests that sound sensitivity based on ‘unaided’ particle motion detection in water will be limited to short ranges, as sound pressure decreases with distance by the inverse square law.

## Sound as a mechanical force – air-borne sound

The physical interaction between sound and the body is different for a terrestrial animal because of the impedance difference between its tissue and the surrounding medium. Thus, a large fraction of sound energy will be reflected by the body of the animal, even when the animal is small compared to the wavelength of the incident sound (i.e., ka<1). Additionally, unlike aquatic organisms, terrestrial organisms are not suspended in the surrounding medium with neutral buoyancy and so friction against the substrate will limit the induced movement. Morse (1941, reprinted 1986) developed an analytical model for the sound-induced movement of a small cylindrical object in air (see appendix). When ka <<1, the model predicts that the induced velocity is largely independent of frequency and can be approximated by 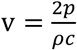 and acceleration 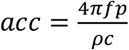 (Morse, 1986, correcting for a 2π error in the derivation; see appendix) resulting in a vibration velocity of v=5.3 µm*s^-1^Pa^-1^ for an object density of 1060 kg/m^3^ and sound velocity 343 m/s. Sensitive eardrums, for example those of mammals, vibrate at approximately 1000 µm*s^-1^Pa^-1^ (Christensen-Dalsgaard and Manley 2013), so if the whole-body vibration velocity was conveyed effectively to the inner ear the sensitivity of a non-tympanic ear would be reduced approximately by a factor 200, or 46 dB compared to the sensitivity of a tympanic ear. The sound-induced velocity declines with increasing *ka*, and this approximation is not valid for animals with *ka*>1. Note, that the induced velocity for equal sound pressures in air and water results in much larger vibrations in air (see box 1); however, the model does not include effects of friction or the mechanical parameters (e.g., elasticity etc.) of the object.

Since the model is an approximation that only holds for low *ka*, we have tested the predictions of this analytical model compared to a simple finite-element model generated using COMSOL Multiphysics software (see details of the model in appendix 2). The model object is a small cylinder with the same density as muscle (1060 kg/m^3^), ensonified by a plane sound wave with a sound pressure of 1 Pa. We modeled the responses of different sized cylinders with similar aspect ratios (1.5:1 length:diameter), spheres and ellipsoids, ranging from the sizes of very small frogs (head diameter 0.01m, length 0.015 m) to large animals (head diameter 0.2 m, length 0.3m). Fig. 1A shows the result of these model responses for different sizes of cylinders. The induced vibration velocities are maximal in the direction of wave propagation and have a constant amplitude of approximately 5 µm/s at low frequencies. The decline in response depends on the size of the object, and when the response is plotted as a function of *ka*, the curves for different object sizes are almost identical (fig. 1B). Thus, the vibrational response to an impinging sound pressure wave depends only on *ka*. At a *ka* of 1, velocity declines by 1.6 dB compared to the maximal amplitude, and a 3-dB decline is found at approximately *ka*=1.2.

**Figure 1.**
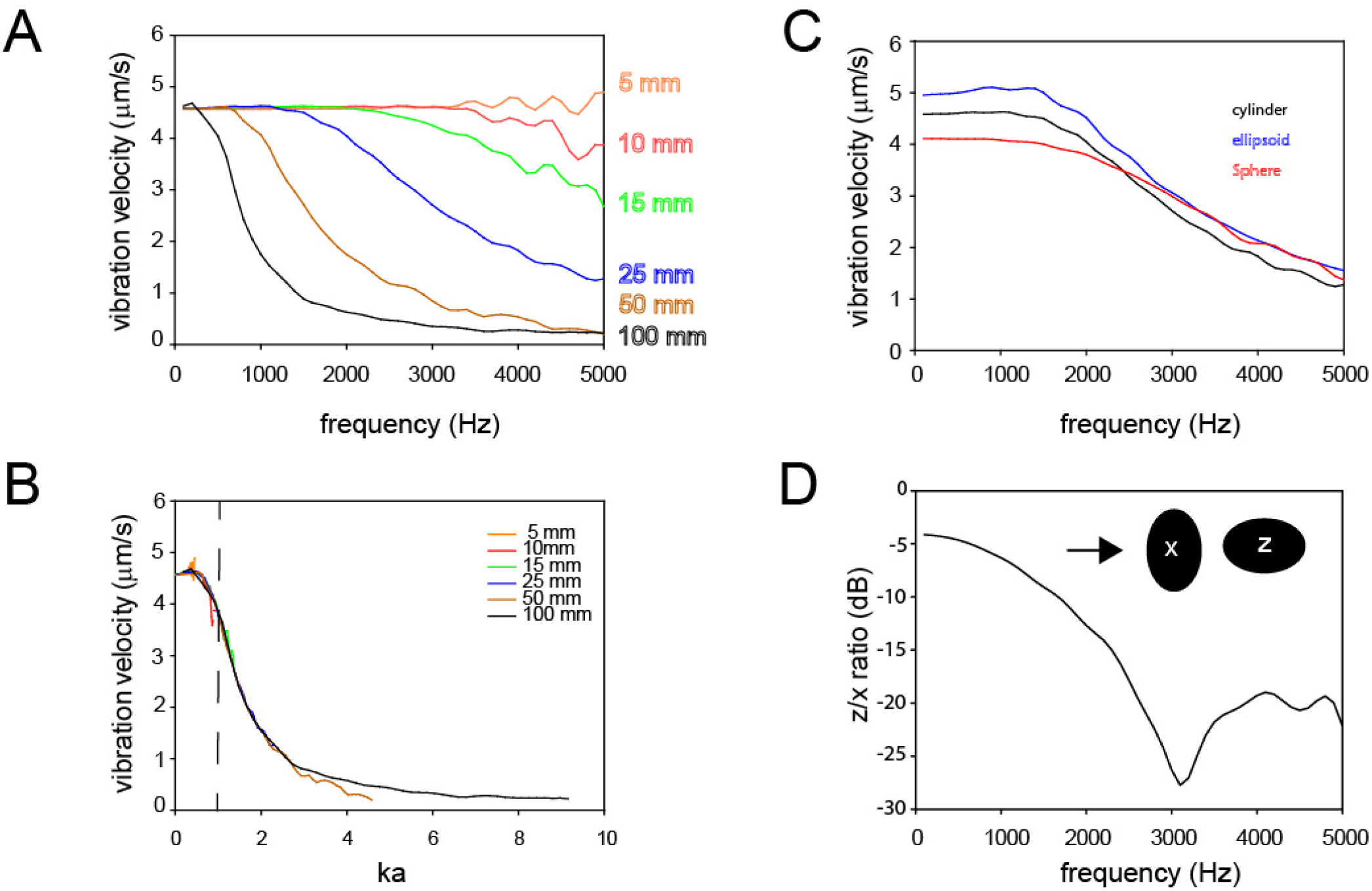
A, B: Finite element model (blue line) of the induced motion of a cylinder of different sizes translated in a sound field (1 Pa), in B shown as a function of ka. C: Model of sound-induced motion of a cylinder, ellipsoid and sphere with the same radius (25 mm).D: ratio of vibration in ellipsoid with sound stimulation perpendicular to the longitudinal axis (X) or parallel to the long axis (Z)

The translation response also depends on object shape. This is shown in fig 1C by comparing the translation response of cylinders, spheres and ellipsoids with similar radius 0.025 m. The three shapes show a similar decline with frequency, but the low-frequency plateau is lower for the sphere model than for the two other models. However, all fall in the range of 4-5 µm*s^-1^Pa^-1^.

Finally, as shown in fig. 1D, the response is direction-dependent, in which greater responses are elicited in cylinders for which the longitudinal axis is perpendicular to the direction of the propagating sound wave.

From these finite element models, we predict that this mode of acoustic stimulation could be sufficient to generate a physiological response in the inner ear of a small-bodied animal in air. If we analogize an animal’s head to a simple cylinder, we may expect that aerial sound pressure at frequencies below 3kHz will be sufficient generate comparable vibration velocities among animals of head sizes smaller than 15 mm. This response drops off rapidly for larger head sizes (25 mm and larger), such that only very low frequencies (0.01-1 kHz) are capable of inducing body vibrations. We additionally predict that this mode of sound energy transduction will be most efficient for sound waves that propagate perpendicular to the longitudinal plane of the animal’s head, or along the inter-aural axis

## Experimental data

The analytical model predicts a simple relationship between sound and induced body vibrations, with a baseline vibration response of approximately 5 µm/s/Pa at low *ka*. It can be compared to experimental data, where auditory sensitivity has been measured either together with measurements of vibrations of the head of the animal or with vibration sensitivity data. The fundamental hypothesis is that if sensitivity to sound pressure can be explained by sensitivity to sound-induced vibrations, sound thresholds recalculated as head vibration thresholds should higher than or comparable to whole-body vibration detection thresholds. Using the analytical model, vibration velocities (in dB re 1 mm/s) are calculated from the sound thresholds (in dB re 20 µPa) simply by subtracting 140 dB.

### Salamanders

Salamanders have highly reduced middle ears that lack the tympanum, middle ear cavity, and Eustachian tubes which, in frogs, connect the two ears through the mouth cavity. Acoustic energy enters the oval window of the salamander inner ear via two middle ear elements, the ossified stapes (columella) and the cartilaginous operculum that articulates with the stapes and has a muscular connection to the pectoral girdle (Kingsbury and Reed, 1908; Monath, 1965; Smith, 1968). In a recent paper on salamander hearing (Capshaw et al., 2020), sound and vibration thresholds were studied using auditory brainstem responses in eight species: two ambystomatid species: *Ambystoma opacum* and *A. tigrinum*, and six plethodontid (lungless) species: *Desmognathus fuscus sp*., *Eurycea cirrigera, E. lucifuga, Gyrinophilus porphyriticus, Plethodon cinereus*, and *Plethodon glutinosus*. Additionally, sound induced vibrations of the heads of salamanders were measured using laser vibrometry. Threshold sensitivities to air-borne sound ranged from 57 to 85 dB SPL, and the most sensitive frequencies ranged from 100-250 Hz. For each species, thresholds to vibrations were almost uniform (in terms of acceleration) up to approximately 300 Hz but varied across species from -55 to - 35 dB re 1 m/s^2^.

Using the laser vibrometry data, sound thresholds could be referred to the sound-induced head vibration and recalculated as vibration thresholds (Fig. 2). Because all species were relatively small animals (largest head radius 1 cm), and their auditory sensitivity is largely confined to frequencies below 400 Hz, we estimate that the *ka* for salamanders is <0.07 and we show averaged data from all species in fig. 2. The measured head vibrations (blue curve) at sound thresholds are comparable to the vibration thresholds (black curve). The analytical model head vibrations predicted from the sound thresholds are approximately 10 dB lower than the head vibrations (red curve).

**Figure 2.**
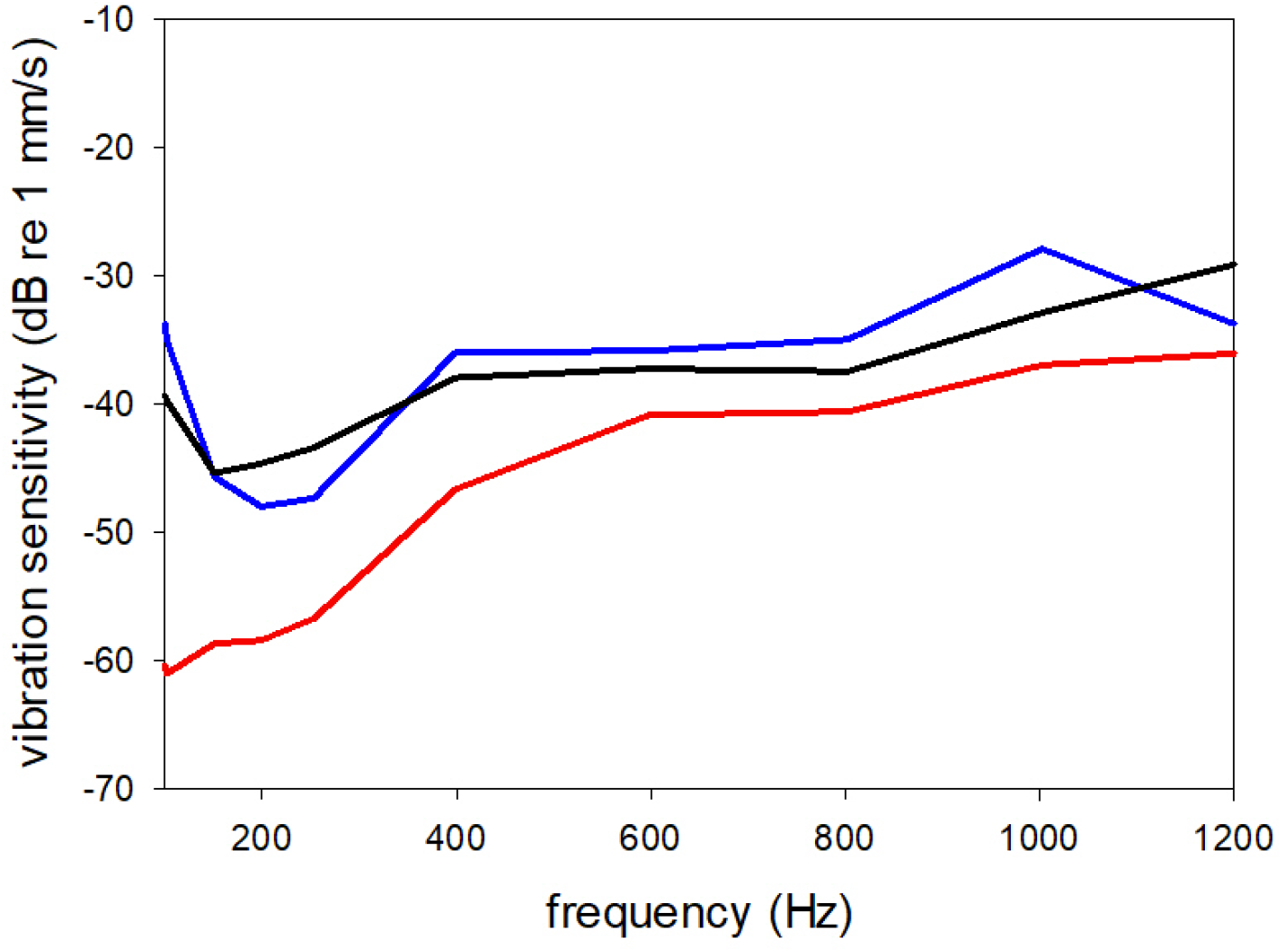
Averaged data from seven salamander species showing thresholds to sound-induced head vibrations measured at the sound thresholds with laser vibrometry (blue curve), compared to seismic vibration thresholds (black curve) and analytical model prediction of induced head vibrations based on the sound thresholds (red curve). Data from Capshaw et al. (2020), J Exp Biol.

A partial explanation of this discrepancy is that the model does not include friction between the animal and substrate, which must limit the movement of the head. Despite the observed discrepancy, this model can be used to estimate the amplitude of accelerations induced in the salamander by air-borne sound. Comparisons of model data to sensitivity thresholds can be used to approximate the efficacy of this extratympanic mechanism for sound detection to evoke a response in the auditory system.

### Earless frogs

Earless frogs are highly diverse in the degree of reduction to their middle ears; at minimum, an earless species lacks tympana, but may also lack the tympanic annulus and stapes (Pereyra et al., 2016). Despite lacking some or all components of the tympanic middle ear, many earless frogs vocalize and presumably detect these acoustic signals via extratympanic pathways (Boistel et al., 2011; Lindquist et al., 1998; Womack et al., 2018). In a study of ten eared and earless bufonid species, Womack et al. (2017) presented data on sensitivity to sound and vibration stimuli. Sound thresholds recorded in eared toads are up to 20 dB more sensitive than the earless toads for frequencies above 900 Hz; however, vibration sensitivity (figure 3) and low frequency (<300 Hz) sound sensitivity is similar among eared (*Rhinella tacana, R. alata, R. leptoscelis, R. spinulosa, R. horribilis*, and *Rhaebo haematiticus*) and earless (*R. arborescandens, R. festae, R. yunga* and *Osornophryne guacamayo*) toads studied. Above approximately 300 Hz eared toads are more slightly more sensitive than the earless, probably reflecting increased stimulation via the tympanic pathway at higher frequencies, and the difference in sensitivity increases with frequency. Womack et al. did not measure sound-induced vibrations in their animals but analytical model calculations of the induced vibrations at the sound thresholds (red curves in fig. 3) results in a prediction of sound-induced vibration thresholds that are lower than the measured vibration thresholds (black curves in fig. 3). This may be caused by the direction of stimulation (dorso-ventral for vibration-stimulation, lateral incidence for sound), but may also suggest that sound is received via alternative and more sensitive pathways than simple translation.

**Figure 3.**
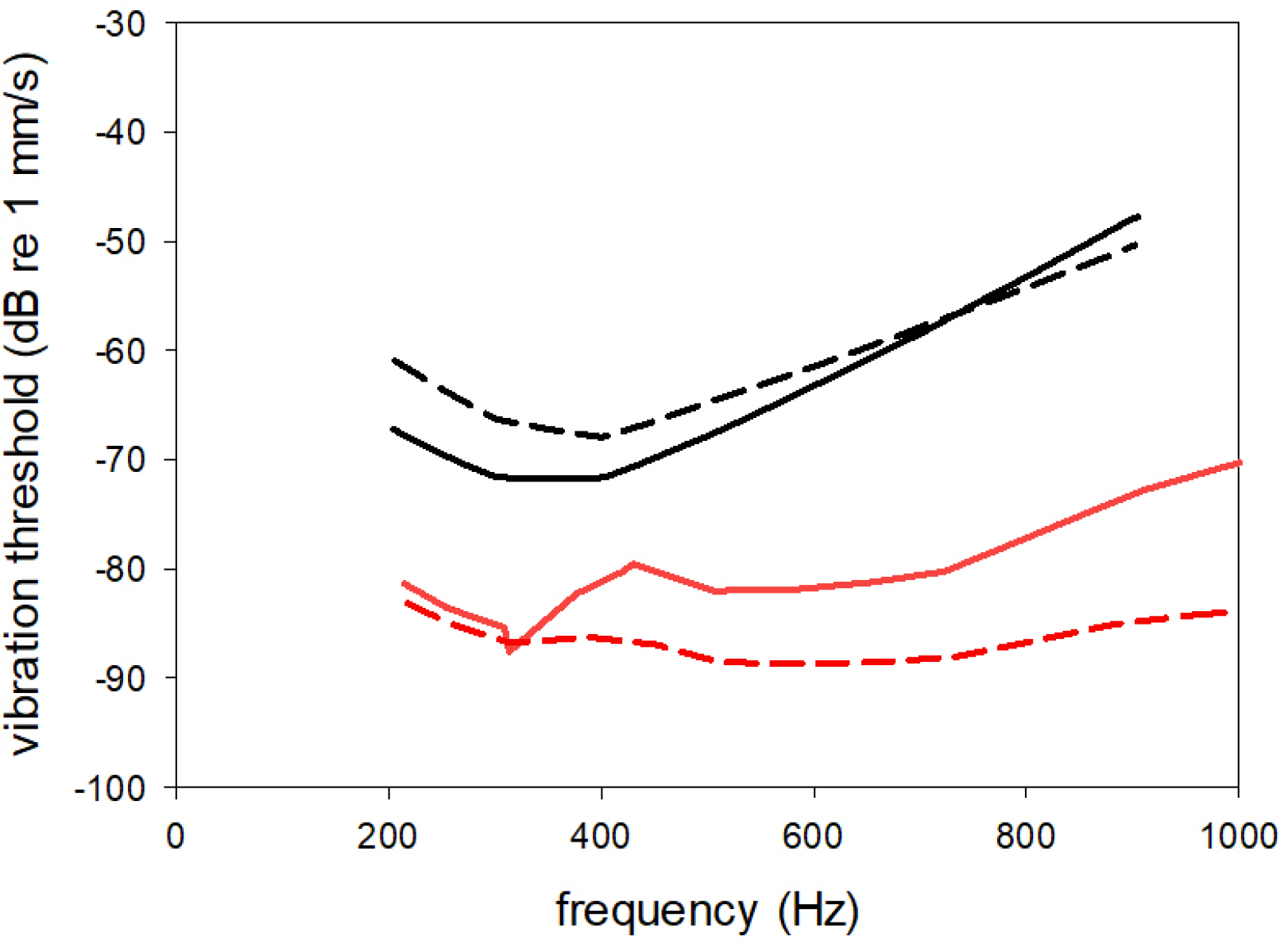
Average vibration thresholds (to dorso-ventral vibration) of earless (black line, data from 4 species) and eared (black dashed line, data from 6 species) toads. The model curves show vibration sensitivity predicted from the thresholds to airborne sound, assuming the sound translation model described in the text (red line: earless, red dashed line: eared). From Womack et al (2017), redrawn.

### Eared frogs

Eared, terrestrial frog species have well-developed tympanic middle ears with flexible tympana coupled to the inner ear via the extrastapes and stapes, in addition to the operculum which articulates with the stapes in the oval window (Mason et al., 2015). Additionally, the middle ear cavities of the two ears are connected to the mouth cavity via the Eustachian tubes. Therefore, the inner ear of tympanate frogs may be stimulated by many different pathways for acoustic energy, including directly though the tympanum or the opercularis system or indirectly through the lung-ear pathway (Ehret et al., 1990; Ehret et al., 1994; Lee et al., 2021; Mason, 2007; Narins et al., 1988). Comparison of sound and vibration sensitivity of single auditory fibers in the eared frog *Rana temporaria* showed that all low-frequency fibers had a dual sensitivity to sound and dorso-ventral vibrations.

The average relative vibration sensitivity, measured as the sensitivity difference between vibration stimulation vibration velocities and sound stimulation particle velocities (fig 4), varied systematically from 42 dB at 100 Hz (i.e., sound particle velocities were 42 dB less efficient) to 25 dB at 400 Hz, probably showing increased sensitivity for tympanic vibrations with increased frequency (Christensen-Dalsgaard and Jørgensen 1996). The analytical model predicts a relative vibration sensitivity of 54 dB (the ratio between sound particle velocity – 2.5 mm/s/Pa – and induced velocity of 5 µm/s/pa), if sound and vibration stimulation were completely equivalent. Thus, sound stimulation was 12 dB more efficient than predicted (and probably even more efficient, since the model does not account for friction), even at 100 Hz. One factor can be that the direction of vibration stimulation (dorso-ventral) maybe is less efficient than the lateral stimulation generated by free-field sound, but alternatively, differential motion of the columella may be larger with sound and vibration stimulation.

**Figure 4.**
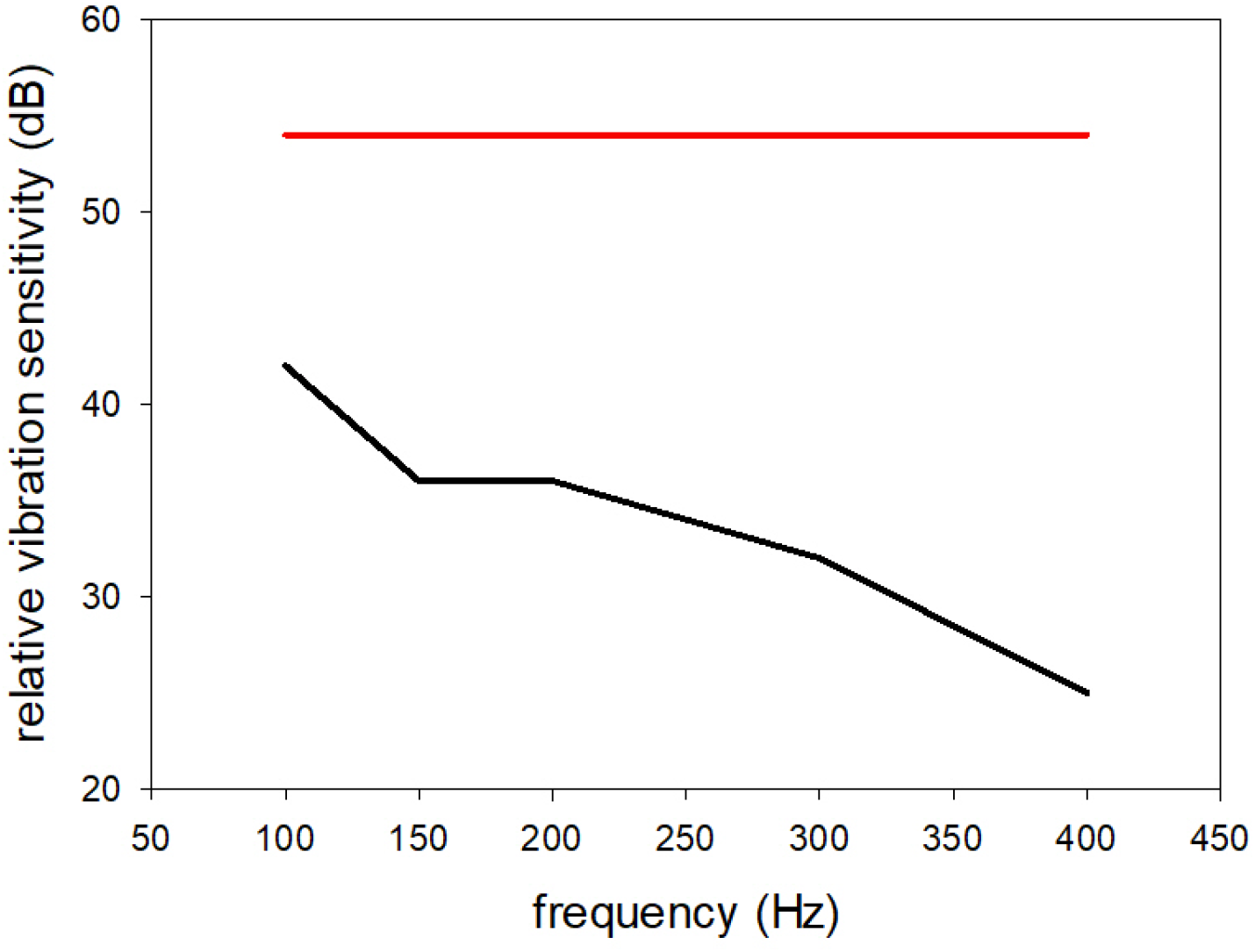
Average vibration sensitivity in single auditory nerve fibers in the grassfrog (<u>Rana temporaria</u>) measured as the difference between sensitivity to sound and dorso-ventral vibration stimulation. (average from 106 fibers). The relative vibration sensitivity predicted from the analytical model is shown as the red curve.

### Snakes

The middle ear of snakes is modified compared to other terrestrial vertebrates as the columella is connected via the quadrate to the lower jaw, and the other tympanic structures have been lost. This is probably an adaptation to detect seismic vibrations. Christensen et al. (2012) studied auditory brainstem responses in the royal python, *Python regius*, to both sound and vibration stimuli and also measured the sound induced head vibrations (Fig. 4). Sensitivities to sound and vibration are most similar from 100-300 Hz. The lowest threshold for sound was 78 dB SPL measured at 160 Hz, corresponding to a model velocity of -62 dB re 1 mm/s. At 500 Hz, the sound threshold level increases to 92 dB SPL, corresponding to a model acceleration of -48 dB re 1 mm/s. Interestingly, analytical thresholds predicted by the model for frequencies above 400 Hz are greater than the sound-induced head vibration thresholds, suggesting a more efficient stimulus pathway for sound at these frequencies.

### Lungfish

The lungfish ear is somewhat typical of an aquatic vertebrate and hearing is mediated by the otolithic organs of the inner ear that detect underwater sound as particle motion and pressure (Christensen et al., 2015). Lungfish hearing in air is rather insensitive and restricted to low frequencies, most likely because it is limited by the frequency response of the otolithic hearing organs. Christensen et al. (2015) measured hearing sensitivity of the African lungfish (*Protopterus annectens*) in air using auditory brainstem responses (Fig. 5). The lowest detection threshold for air-borne sound was 86 dB re 20 µPa, measured at 80 Hz, and the threshold increased to approximately 100 dB re 20 µPa. at 200 Hz. For lungfish, the head radius is approximately 0.05 m, so ka<1 for frequencies below approximately 1 kHz. The analytical model calculation predicts a vibration amplitude of -54 dB re 1 mm/s at 80 Hz and -40 dB at 200 Hz (fig. 5, red line). Again, the model predicts thresholds that are lower than what is actually measured, because the predicted head vibrations are greater than the measured vibrations.

**Figure 5.**
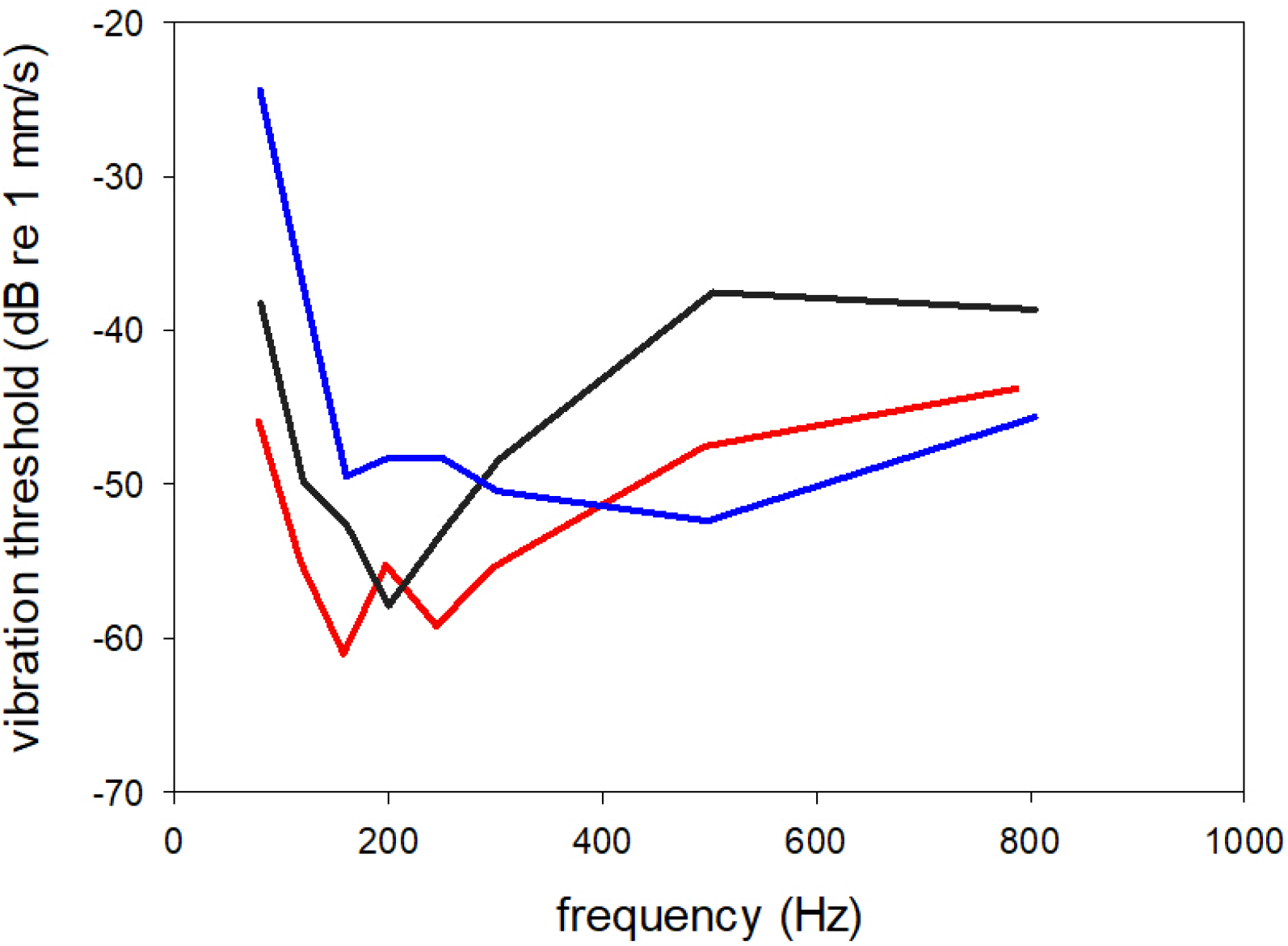
Average sound and vibration thresholds of royal pythons, Python regius. Sound thresholds are referred to the measured sound-induced head vibrations (blue). The red curve shows the model-predicted induced vibrations, and the black curve is experimentally measured vibration thresholds. From Christensen et al. 2012, redrawn.

**Figure 6.**
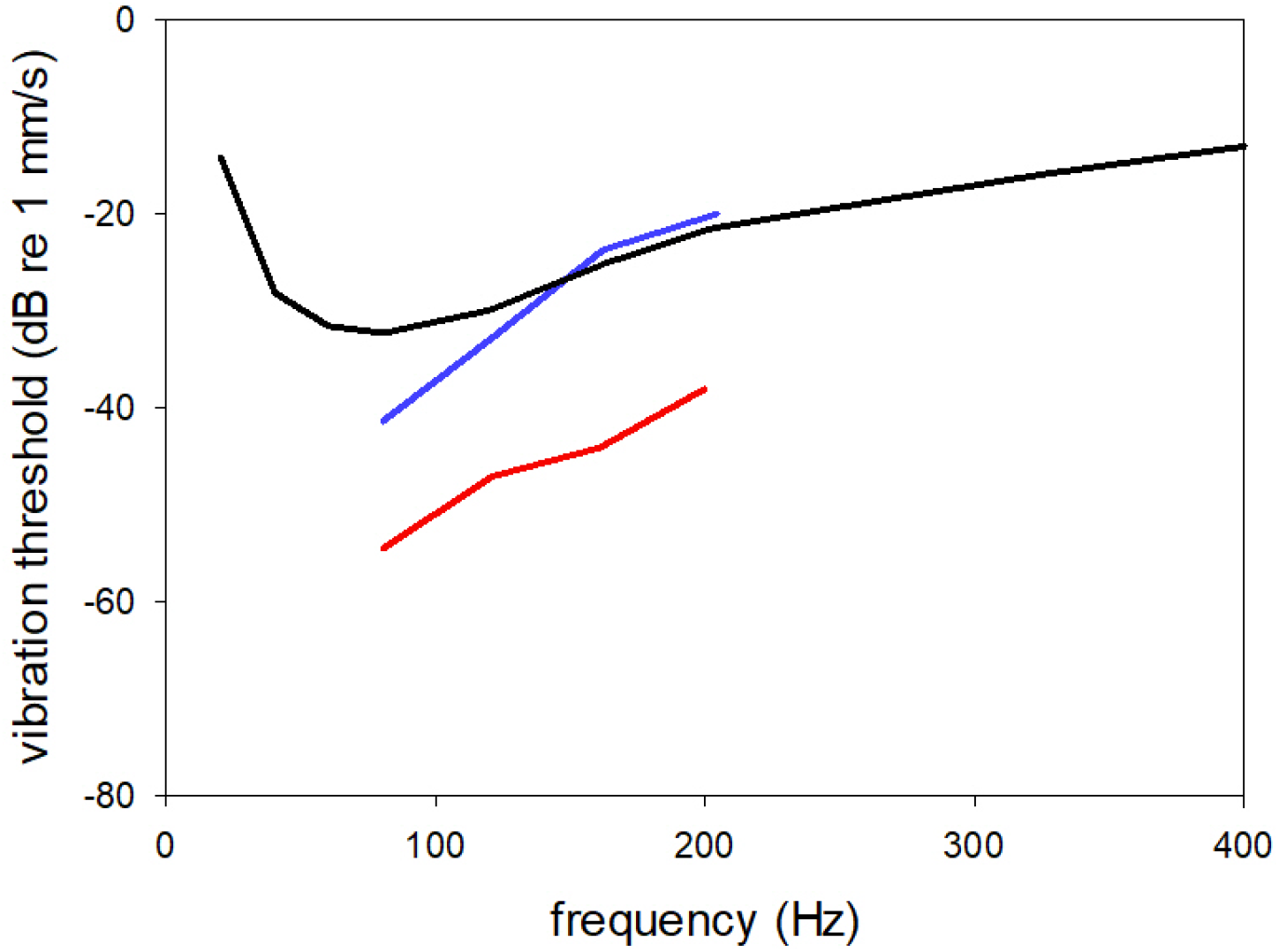
Thresholds to air-borne sound and vibration in African lungfish Protopterus annectens. The sound thresholds (blue) are referred to the sound-induced head vibrations for comparison with the vibration thresholds (black). The model head vibrations calculated from the sound thresholds is shown as the red curve. From Christensen et al 2015, redrawn.

## Discussion

The experimental data reviewed here show similar ranges of sound sensitivities among all atympanate animals studied. Auditory sensitivity is restricted to low frequencies (defined by *ka*, the acoustical size of the animal) and can in most cases be explained by sensitivity to sound-induced vibrations of the head of the animal. We used a theoretical model to predict the ability of sound to induce translational movement in these atympanate animals, approximated as a simplified cylinder with the viscoelastic properties of muscle. In all cases the analytical model predicted 10-15 dB larger sound-induced vibrations (hence lower sound thresholds) than have been measured, so even though this simple bone conduction mechanism for sound detection seems to generate sufficient amplitude vibrations in atympanate animals to explain the auditory sensitivity the induced vibrations are probably at least 10 dB lower than modelled, so closer to 1 µm/s/Pa.

The most likely explanation for the discrepancy between modelled and actual vibrations is that the model does not account for friction, which must be important, perhaps especially at high frequencies. Also, the motion of head and body is likely not uniform, and the head will move relative to the body of the animal. Also, cranial vibrations likely will also contribute to non-tympanic hearing, as is the case in humans (Stenfelt 2013), where structural vibrations dominate bone-conduction sensitivity above approximately 400 Hz. However, such vibrations will be strongly dependent on the anatomy and material properties of the head and might contribute at lower *ka* in other animals. Resonances in air cavities could also contribute to the vibrations of the head, as suggested by Boistel et al. (2011).

However, our findings provide support for a general mechanism for sound reception without any specialized middle ear apparatus. This mechanism would have been functional in the earliest tetrapods that may have had an ear configuration similar to the lungfish. Interestingly, this mechanism for sound-induced head translation is also proposed for human bone conduction frequencies below 400 Hz (i.e. *ka* < 0.62 assuming a head radius of 0.085m) (Stenfelt, 2013). A sound-induced vibration of the human head of 3 µm/s/Pa, as proposed by von Bekesý (1960) fit the measured human bone conduction thresholds at low frequencies reasonably well (see Sørensen et al. 2022). Additionally, the comparison of sensitivity of eared and earless bufonids by Womack et al (2017) shows that, for low-frequency hearing, earless species are not really at a disadvantage relative to tympanate species, indicated by nearly equivalent thresholds to sounds below 900 Hz. Thus, this bone conduction mechanism for hearing may explain the relaxed selection pressures for functioning middle ears in the anurans. In anuran amphibians, the structures of the tympanic middle ear form relatively late during development, and reduction or loss of the tympanic middle ear may be correlated to developmental heterochronies associated with paedomorphosis and/or miniaturization of body size (Hetherington, 1992; Pereyra et al., 2016). Especially in connection with miniaturization, an allometric downsizing of the tympanum would increase its stiffness and therefore reduce sensitivity to air-borne sound and may further relax the selection pressure for functional middle ears.

It remains a question how the whole-body vibration is stimulating the inner ear, and some of the mechanisms reported for human bone conduction, such as inertial movement of the middle ear ossicles or vibrations traveling through the cranial bone to the inner ear fluids, are not likely to function for simple translation by sound. In fish, sound reception is facilitated by the inertial vibrations of the otoliths, but the otoliths of tetrapods, at least in amphibians, are only sensitive in a restricted frequency range (Christensen-Dalsgaard and Jørgensen, 1988). Alternatively, inertia of the inner ear fluids has been proposed as a the dominant mechanism for human bone conduction (Stenfelt, 2015) and may function in other animals as well (see review in Capshaw et al. 2022). Fluid inertia will produce relative motion of the inner ear fluids and the sensory epithelia. Sensitivity to fluid inertia would depend on the size and orientation of pressure release windows in the inner ear but is a very general mechanism that would function in a variety of tetrapods.

## Conclusion

The sound-induced vibrations for animals with small acoustical size, indicated by low ka, provides a baseline auditory sensitivity that explains terrestrial hearing abilities observed in diverse animals without specialized middle-ear structures. This hearing mechanism is effective at low frequencies where the eardrum in small animals is less responsive. Before the origin of the tympanic middle ear, sound-induced vibrations likely was the mechanism for stimulation of the ear in early tetrapods and could potentially even provide directional information (Capshaw et al. 2022). This may have provided an important foundation for pressure sensitivity on land, until a series of modifications leading to a functional middle ear increased sensitivity to airborne sound (Christensen-Dalsgaard and Carr 2007). These modifications are proposed in a three-step model of middle ear evolution by Christensen-Dalsgaard and Manley (2014): a reduction of otolith covering in the sensory macula, development of movable elements such as columella or operculum and finally formation of columella-tympanum connection.

## List of Symbols

c: sound velocity
f: frequency
λ: wavelength
k: wave number
a: radius
p: sound pressure
Z: specific impedance
v_p_: particle velocity
v: vibration velocity
acc: acceleration
ρ(rho): density

## Appendix. 1.

Morse: Force on cylinder (Morse 1981, p 352)

This calculates the acoustical force on an object – a small cylinder of unit length. The approximations are for low ka, i.e. objects that are small compared to the wavelength of sound. It is usually stated that such objects are ‘transparent’ to sound, but locally, since the object motion must be equal to the particle velocity of the medium bordering the object, a part of the sound wave is reflected. Therefore, the equations below incorporate the (frequency dependent) reflection coefficients.

The force is pressure times area, where

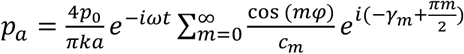 a is the radius. and 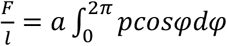

C_m_, γ_m_ are the reflection coefficient amplitudes and phases of the cylinder (Morse 26.6, p.301)

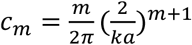.

At <u>low ka</u>, 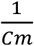 coefficients are small when m>1, and a useful approximation is therefore to use only the two first elements (c_0_ and c_1_)

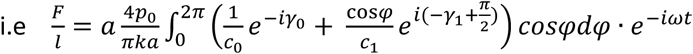

The first (c_0_) term in the integral is constant in φ, so the integral is zero (integrating a cos function over a whole cycle), resulting in the integral

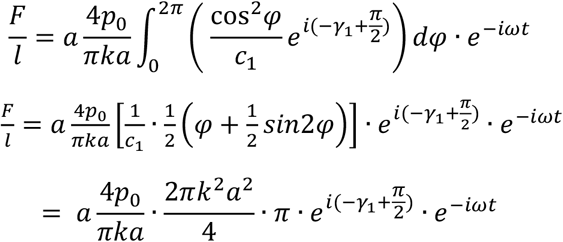

The amplitude is 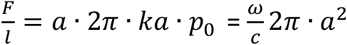

Acceleration, 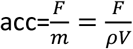 where m is mass, ρ density and V volume, *V* = π*a*^2^*l*

Thus, 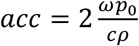, velocity 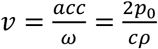, and displacement 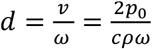

Note, that the formula stated by von Békésy for a sphere at low ka (1960, p. 171) is 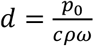, a factor 2 lower.

## Appendix. 2.

Comsol-model.

The finite-element models of ensonified objects were made in Comsol Multiphysics, v. 5.6. We used simple geometric shapes – cylinders, spheres and ellipsoids, with varying radii and (for cylinders and ellipsoids) the same length/diameter ratio (1.5). The model simulated the shapes suspended in air (frictionless) and ensonified by plane sound waves from different directions (parallel and perpendicular to the long axis).

The meshing was triangular, using a physics-controlled mesh with ‘finer’ resolution.

The material of the shapes had a density like muscle tissue (1060 kg/m^3^), but to avoid internal vibrations in the structure we made the material unrealistically stiff (Youngs modulus 45 * 10^9^ Pa), more than 1000 times higher than normal tissue.

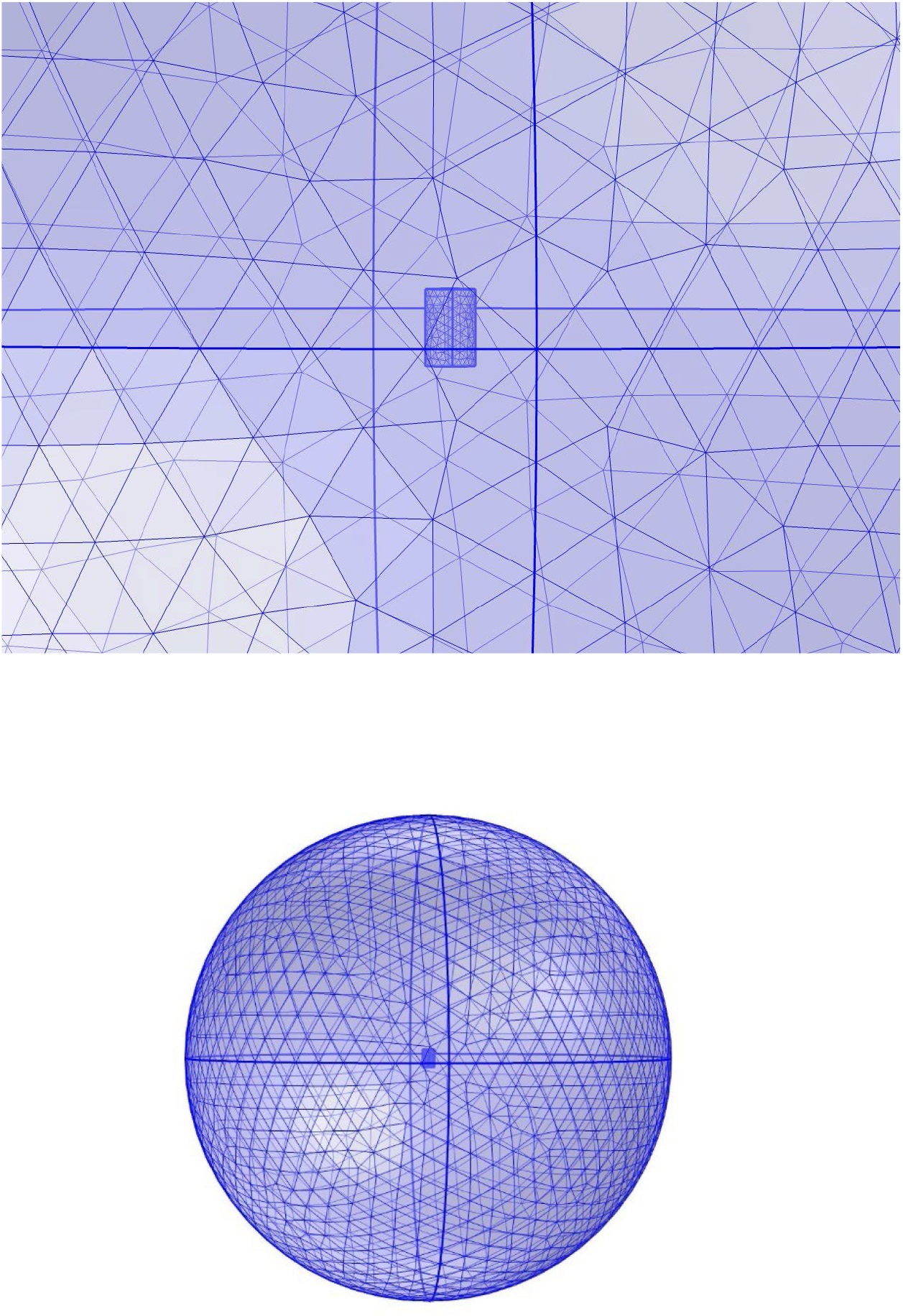

## Notes

### Competing Interest Statement

The authors have declared no competing interest.

